# Performance of pathogen identification and resistance gene expression tests using ASTar® remnant bacterial suspension in Gram-negative contrived positive blood cultures

**DOI:** 10.64898/2026.07.01.735973

**Authors:** Vikas Gupta, Michelle Myers, Ida Niklasson, Stina Vincentsson, Elin Ring, Oliver Mainwaring, Natalie Brown, Jan Grawé

## Abstract

**Introduction:** Rapid pathogen identification, resistance detection, and susceptibility profiling improve antimicrobial prescribing and associated outcomes, but fragmented workflows lead to inefficiencies and are costly. We evaluated a research-use-only (RUO) approach using ASTar® remnant bacterial suspension from routine AST for MALDI-TOF MS pathogen identification and Lateral Flow Assay (LFA)-based detection of targeted resistance mechanisms.

**Methods:** Gram-negative (GN) bacterial strains from reference and curated resistance collections (CDC1, ARLG2, ATCC3) [n=119] were contrived into blood culture bottles and processed in the ASTar System using the ASTar BC G-Kit (Q-linea AB, Sweden). Under RUO conditions, remnant bacterial suspensions were collected ~1-2 h after ASTar run initiation and analyzed using NG Test CTX-M Multi, NG Test CARBA-5, NG Test Acineto-5 RUO, and MALDI-TOF MS.

**Results:** Mean (± SD) remnant suspension volume was 2722 μL (± 300 μL). All samples yielded high-confidence MALDI-TOF MS scores (>2.0), with five initially scoring <2.0 and resolving on repeat testing. LFA results showed full agreement with reference isolates for *blaCTX-M* positive/negative (30/30) and with 60 or 61 target carbapenemase-positive/negative isolates. Testing of a subset of samples to mimic reflex workflows with ASTar phenotypic results did not affect LFA performance (n=26; 23 Enterobacterales, 3 *P. aeruginosa* and 9 *A. baumannii)*. Cost savings can be realised versus commercial multiplex PCR.

**Conclusion:** This integrated approach of ~6 h rapid phenotypic AST with same-run identification and resistance detection (1-2 h from instrument start) or reflex testing upon availability of ASTar results may support earlier susceptibility results and offer cost savings to current workflows.

## Introduction

Bloodstream infections (BSI) represent the most severe form of bacterial infections encountered in clinical practice and are often a precursor to sepsis, a life-threatening condition defined by organ dysfunction resulting from a dysregulated host response to infection. Sepsis is a major global health burden, affecting more than 166 million people every year and accounting for approximately 21.4 million deaths worldwide. The incidence of sepsis continues to rise, in part due to increasing rates of global antimicrobial resistance compromising the efficacy of antimicrobial treatments^1^.

Optimal management of patients with BSIs and sepsis relies on rapid identification of the causative pathogen and its antimicrobial susceptibility profile. Delays in antimicrobial treatment are associated with worsened clinical outcomes, and early administration of appropriate antimicrobial therapy is among the strongest predictors of survival^2^. Clinical guidelines therefore recommend initiation of antimicrobial therapy within one hour of probable or definite sepsis recognition^3^. In practice, this is achieved through empiric broad-spectrum therapy while awaiting definitive microbiological results including full pathogen ID, resistance markers, and antimicrobial susceptibility testing (AST) data.

Conventional workflows for organism ID and AST vary across countries, healthcare systems, and laboratories, but typically follow a common sequence^4^. Following blood culture collection, incubation, and positivity, samples are Gram-stained and subcultured onto appropriate solid media to obtain isolated colonies, followed by standard incubation and colony growth prior to organism ID and susceptibility testing. This process commonly takes 24-72 hours, in the absence of further delays, before actionable AST results are available and tailored patient treatment can begin^5^.

Consequently, recent practice guidelines from the American Society of Microbiology recommend rapid diagnostic tests along with a comprehensive plan to communicate results in order to improve time to target therapy (TTT) (reference 4). Rapid pathogen ID platforms, including molecular assays, can provide organism ID and detect resistance genes; however, these approaches are often associated with higher costs and identify the presence of resistance genes without reliably predicting phenotypic expression.^8^ Lateral Flow Assays (LFAs) can detect expressed resistance markers, serving as a supplementary method that may help bridge genotypic detection with phenotypic expression.

MALDI-TOF MS is widely used for rapid organism ID due to its high accuracy and low per-test cost, in addition to broad organism libraries. However, conventional MALDI workflows require isolated colonies, which can introduce delays from overnight subculture. Direct-from-blood-culture MALDI methods exist but often require additional manual processing steps. Lateral flow assays (LFA) are generally less complex methods that enable rapid detection of specific resistance markers directly from blood culture material. While LFAs do offer speed, simplicity and lower cost compared to commercial syndromic testing they are limited to predefined targets and do not provide comprehensive antimicrobial susceptibility information^8,9^.

ASTar® (Q-linea AB, Sweden) is a rapid phenotypic AST system that can shorten the time to phenotypic susceptibility results by ~36-42 hours (reference 6, 7 add Garrett et al). During routine sample preparation ASTar produces a purified bacterial suspension within its disposable reagent cartridge after approximately one hour of assay start, part of which is discarded as remnant material upon completion of the standard procedure. While the intended use of the ASTar System is limited to AST only and it is neither CE-marked nor FDA-cleared for sample preparation for organism ID or use in LFA, this remnant material generated by the system may provide opportunity for potential research-use-only (RUO) downstream analytical applications, including MALDI-TOF MS and LFAs outside the scope of the system’s approved claims.

In this study, we evaluated the feasibility of using ASTar remnant material for MALDI-TOF MS pathogen identification and LFA-based resistance detection. This work was conducted exclusively under RUO conditions with the aim of assessing the ASTar remnant bacterial suspension and generate foundational data supporting the future development of integrated time-efficient workflows for comprehensive rapid diagnostics in BSIs and sepsis.

### Importance

Rapid, integrated diagnostic workflows are recommended and essential for improving patient outcomes in bloodstream infections, where delays in pathogen identification and resistance detection can compromise timely antimicrobial therapy. Current workflows often rely on multiple platforms, increasing turnaround time and laboratory burden. Additionally, the clinical impact of rapid diagnostic results is dependent on timely interpretation and implementation within antimicrobial stewardship frameworks, which may involve complex decision algorithms.

This study demonstrates that remnant bacterial suspensions generated during routine AST using the ASTar System can be recovered and utilized for rapid MALDI-TOF MS and Lateral Flow Assay-based resistance detection. By enabling pathogen identification and/or resistance detection within ~1-2 h of AST initiation or resistance detection via reflex testing, this approach reduces duplicate sample testing and may enable earlier access to rapid microbiological results.

Furthermore, this approach offers a cost-saving alternative to molecular methods such as multiplex PCR while concurrently maintaining high diagnostic accuracy, supporting more efficient clinical microbiology workflows.

## Materials and methods

### ASTar

The ASTar System (Q-linea AB, Sweden) is a fully automated, rapid phenotypic antimicrobial susceptibility testing (AST) platform that is CE-IVDR certified and FDA 510(k) cleared for use directly from positive blood cultures. The system consists of the ASTar Instrument and the ASTar BC G-Kit. The kit comprises two disposable consumables: a cartridge containing all reagents required for sample preparation and AST processing, and the 336-well disc containing 23 antimicrobial agents on the current FDA-cleared panel and 24 on the current CE-IVDR-marked panel across a broad range of concentrations.

Following confirmation of monomicrobial Gram-negative blood culture positivity, 1 mL of positive blood culture material is directly loaded into the ASTar cartridge, where the sample undergoes automated concentration determination and dilution steps to generate a final bacterial inoculum aligned with both EUCAST and CLSI guidelines. The ASTar disc is then inoculated, and a remnant bacterial suspension remains within the cartridge. AST is subsequently carried out across the antimicrobial panel, and bacterial growth is monitored using time-lapse microscopic imaging in combination with proprietary algorithms, enabling determination of minimum inhibitory concentrations (MICs) and corresponding S/I/R categorizations. Total system sample preparation time is approximately one h, and comprehensive AST reports are delivered approximately six hours from the start of the ASTar run.

### Workflow

Well-characterized Gram-negative strains with available pathogen identity and genotypic results (CDC1, ARLG2, ATCC3) were contrived into blood culture bottles. Following blood culture positivity, the samples were loaded into the ASTar system following the manufacturer’s instructions. Under a research-use-only (RUO) protocol, remnant bacterial suspensions were removed from the ASTar cartridge after approximately one hour of assay start and, where applicable, tested using one or more LFAs (NG TEST CTX-M Multi, NG Test CARBA 5, NG Test Acineto-5 RUO; NG Biotech), and rapid ID by MALDI-TOF MS (Bruker Biotyper MALDI System). For each assay preparation, a 600 µL aliquot of remnant suspension was centrifuged for three minutes at 12,000 g. The supernatant was carefully aspirated by pipette to retain an undisturbed dry pellet at the bottom of the centrifuge tube for subsequent applications.

To prepare the pellet for MALDI-TOF MS, the pellet was directly spotted on a MALDI target plate, MALDI matrix was then added directly to the smear and allowed to dry before identification was performed using the Bruker Biotyper MALDI system (IVD library). Repeat MALDI testing was performed for isolates that had a confidence score of <2.0 on the initial MALDI run. A confidence score of ≥ 2.0 was considered as identification of the pathogen, and repeat testing was not necessary for these samples. MALDI scoring was evaluated and compared to the reference bacterial identity.

To prepare the pellet for LFA testing, the pellet was resuspended in 150 uL of the kit-provided extraction buffer. From this, 100 µL was applied to the test cassette and incubated for 15 minutes prior to interpretation. To mimic a reflex workflow following the availability of final phenotypic AST results from ASTar, cartridges from a subset of samples ([n=26], 23 *Enterobacterales*, 3 *P. aeruginosa*) were stored refrigerated for six hours after retrieval from the instrument before performing the pellet preparation and LFA assay. Another nine *A. baumanii* samples were stored refrigerated for six hours after retrieval of the cartridge before testing on an RUO Acineto-5 assay. LFA results for detected resistance mechanisms were compared with the reference bacterial genotype, and positive predictive value (PPV) and negative predictive value (NPV) were calculated.

### Cost Comparison

Incremental additional cost or cost savings were modelled for low-, medium- and high-prevalence settings for carbapenem resistance (Enterobacterales, *P. aeruginosa*/*A. baumannii*) and ESBL production (Enterobacterales), in institutions with and without commercial multiplex systems. Low- and high-cost assumptions, expressed in US dollars (USD, $), are detailed in Table 1. ASTar System costs were excluded because they were assumed to remain constant across all comparisons.

**Table 1.** Cost assumptions for different methods for bacterial identification and resistance markers.

| Cost Assumptions |  |  |
| --- | --- | --- |
| Item | Low | High |
| Syndromic molecular Cost | \$70.00 | \$ 120.00 |
| MALDI Cost | \$ 1.00 | \$ 1.00 |
| LFA Carba 5 Cost | \$ 30.00 | \$ 40.00 |
| LFA CTX-M Cost | \$ 25.00 | \$ 30.00 |

## Results

### Workflow

Preparation of the ASTar remnant bacterial suspension for downstream testing was straightforward and required < 10 minutes hands-on time. The remnant was readily available following standard ASTar AST processing, and both MALDI-TOF MS and LFA runs could be prepared from this remnant material in under ten minutes per sample. (Figure 1).

**Figure 1.**
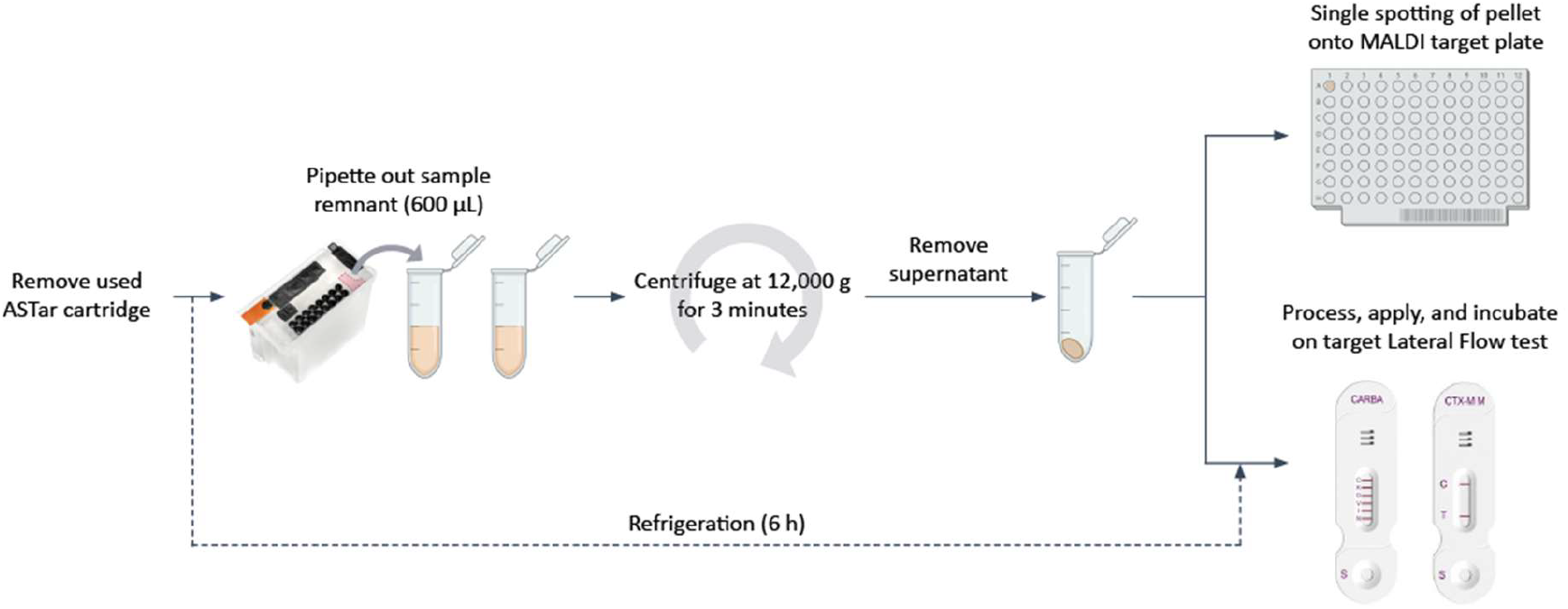
MALDI-TOF MS and LFA preparation from ASTar remnant bacterial suspension.

### MALDI scoring and LFA reference comparison

After cartridge retrieval, the mean (± SD) remnant suspension in each cartridge was 2722 µL (± 300 µL) across 119 Gram-negative samples run on ASTar. Species or species group distribution identified by rapid MALDI was 82% (98/119) *Enterobacterales* [ENT], 15% (18/119) *P. aeruginosa* [PsA]/*A. baumannii [AcB]*, 3% (3/119) *S. maltophilia* (off panel), and 1.7% (2/119) Salmonella spp. (off panel). All 119 samples yielded final high-confidence ID scores >2.0 (Figure 2), including five samples which initially scored <2.0 and re-scored high-confidence on single spotting re-test.

**Figure 2.**
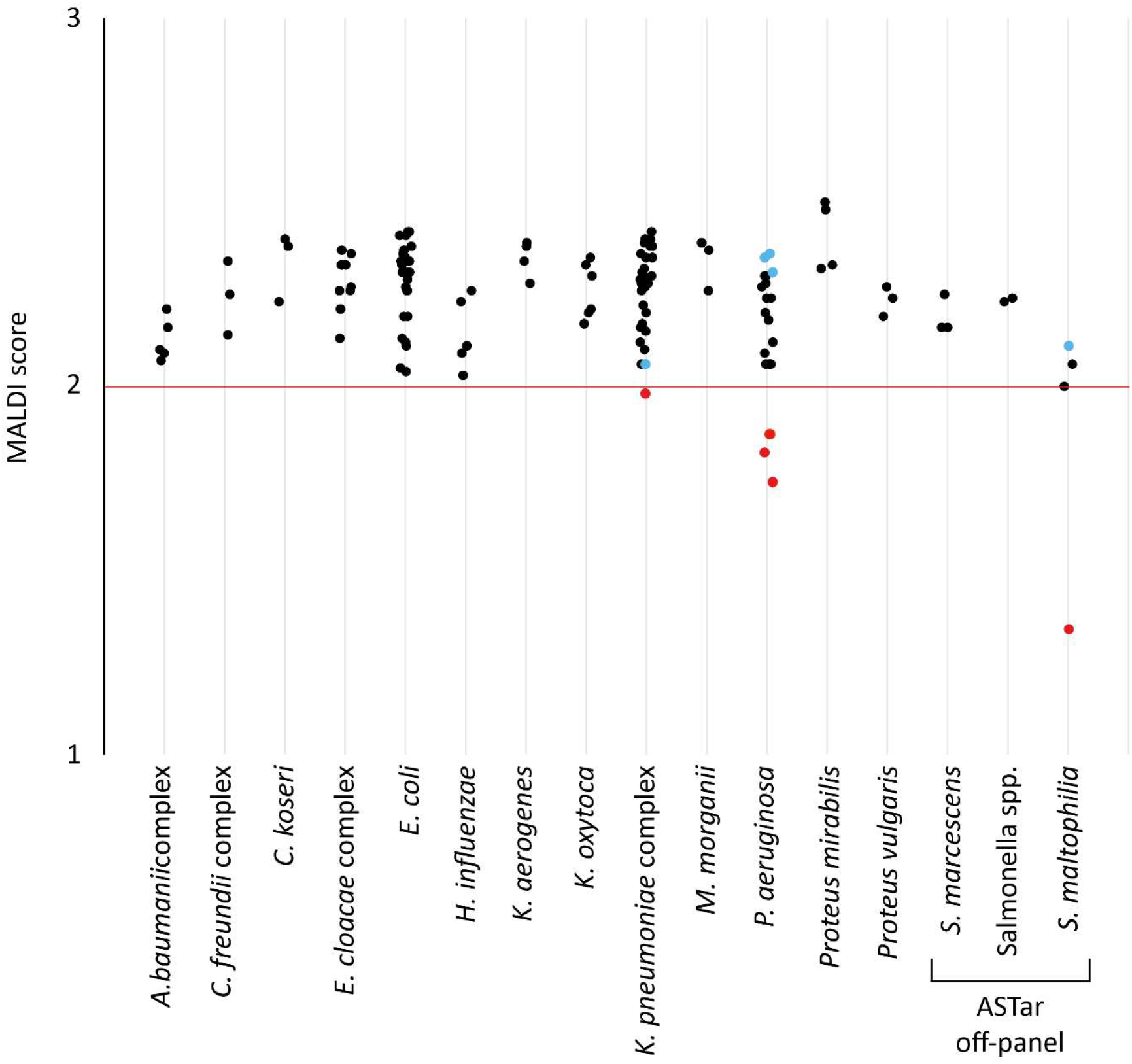
MALDI scores for 119 samples prepared from remnant ASTar output. Red dots: initial score <2.0; Blue dots: score re-run samples which were initially red (<2.0), see methods.

LFA results run after cartridge retrieval (approximately one h after ASTar start) showed full agreement with reference blaCTX-M positive (30/30) and all carbapenemase-positive types, except for one reference OXA-48-positive isolate (60/61) (Table 2). The overall positive predictive value (PPV) was 100% and the overall negative predictive value (NPV) was 96.4%.

**Table 2.** Comparison of LFA testing using ASTar remnant suspension in genotypic confirmed isolates.

| Resistance mechanism | LFA results ASTar Sample Prep, n^ | Expected resistance mechanisms (Reference, N) | % Concordance (n/N) |
| --- | --- | --- | --- |
| **CTX-M Multi Enterobacterales** | **30** | **30** | **100%** |
| CTX-M Group 1 | 13 | 13 | 100% |
| CTX-M Group 2 | 1 | 1 | 100% |
| CTX-M Group 9 | 4 | 4 | 100% |
| Negative CTX-M | 12 | 12 | 100% |
| **CARBA-5 Enterobacterales** | **51** | **52** | **98%** |
| KPC | 17 | 17 | 100% |
| NDM | 11 | 11 | 100% |
| VIM | 3 | 3 | 100% |
| IMP | 4 | 4 | 100% |
| OXA-48 | 4* | 5 | 80% |
| Negative Carba | 12 | 12 | 100% |
| **CARBA-5 *P. aeruginosa*** | **9** | **9** | **100%** |
| KPC | 1 | 1 | 100% |
| VIM | 2 | 2 | 100% |
| IMP | 3 | 3 | 100% |
| Negative carba | 3 | 3 | 100% |

\* 1 isolate initially identified as OXA-48 negative was OXA positive on rerun
^ 26 isolates were stored refrigerated and tested 6 h after retrieval from cartridge with same results

For the subset of 26 samples stored refrigerated for six hours after retrieval of the cartridge before performing the LFA assay (23 *Enterobacterales*, 3 *P. aeruginosa*), reflex LFA results were identical to those from the samples processed directly after cartridge retrieval. Similar results were also obtained for the nine *A. baumannii* samples stored refrigerated for six hours after retrieval of the cartridge before testing on the RUO Acineto-5 assay (Table 3).

**Table 3.**
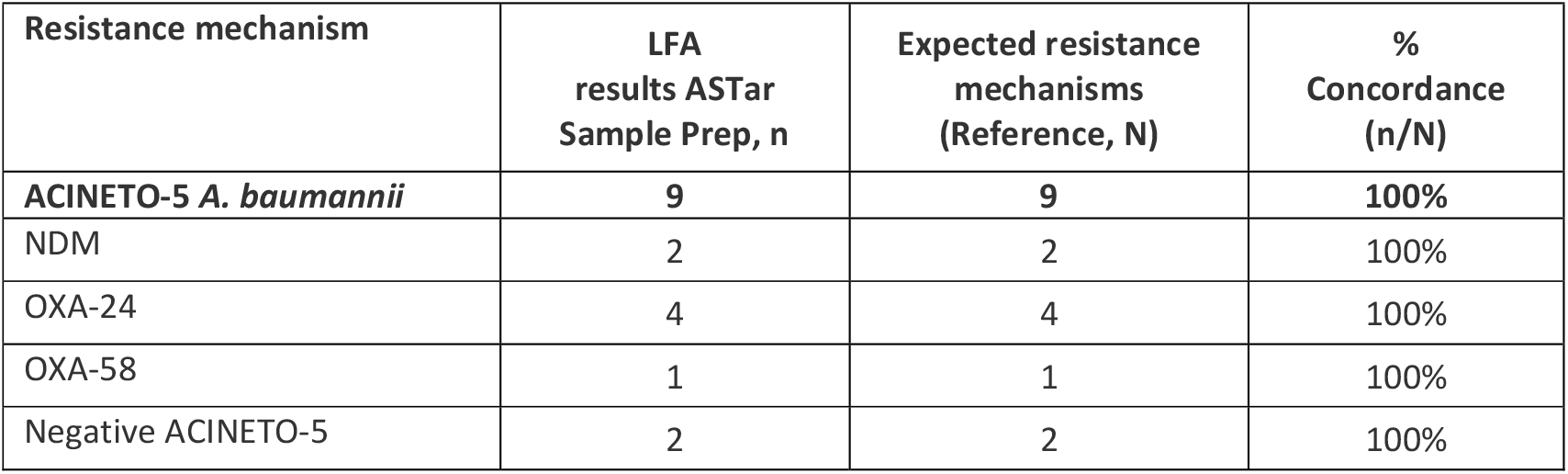
Comparison of LFA testing using ASTar remnant suspension stored refrigerated for 6 h on genotypic confirmed *A. baumannii* isolates.

| Resistance mechanism | LFA results ASTar Sample Prep, n | Expected resistance mechanisms (Reference, N) | % Concordance (n/N) |
| --- | --- | --- | --- |
| **ACINETO-5 *A. baumannii*** | **9** | **9** | **100%** |
| NDM | 2 | 2 | 100% |
| OXA-24 | 4 | 4 | 100% |
| OXA-58 | 1 | 1 | 100% |
| Negative ACINETO-5 | 2 | 2 | 100% |

### Cost comparison using ASTar remnant bacterial sample

The incremental cost of MALDI-TOF MS and reflex LFA testing was evaluated across low-, medium-, and high-resistance settings. Reflex LFA testing was assumed to include CARBA-5 for carbapenem-resistant Enterobacterales, *P. aeruginosa*, and *A. baumannii*, and both CARBA-5/CTX-M assays for carbapenem-resistant Enterobacterales (Table 4). In institutions where syndromic molecular testing is not available, the incremental cost (USD) of MALDI-TOF MS and reflex LFA ranges from $1.39 to $4.15 for Enterobacterales and from $4.60 to $17.00 for *P. aeruginosa/A. baumannii* assuming constant ASTar System cost. In institutions where syndromic molecular testing is available, the incremental cost savings for MALDI-TOF MS and reflex LFA range from $68.62 to $115.85 for Enterobacterales and from $65.40 to $103.00 for *P. aeruginosa/A. baumannii* assuming constant ASTar System cost.

**Table 4.** Cost comparison by system and system combination^.

| **AMR Range (Carb NS/ESBL + ENT)** |  |  |  |
| --- | --- | --- | --- |
|  | **Low** | **Med** | **High** |
| Suggested Carb NS ENT (%) | 0.7% | 2.0% | 4.5% |
| Suggested ESBL (%) | 5.0% | 12.0% | 20.0% |
| Molecular Low Cost | \$ 70.00 | \$70.00 | \$70.00 |
| Molecular High Cost | \$120.00 | \$120.00 | \$120.00 |
| **Cost of Carb NS ENT (Carba 5) + ESBL + (CTX-M)** |  |  |  |
| ASTar Rapid MALDI + Reflex LFA w/o Syndromic Molecular (Low) | \$1.39 | \$ 2.10 | \$ 3.48 |
| ASTar Rapid MALDI + Reflex LFA High Cost (High) | \$1.49 | \$ 2.40 | \$ 4.15 |
| Cost Savings Over Syndromic Molecular (Low) | \$68.62 | \$ 67.90 | \$ 66.53 |
| Cost Savings Over Syndromic Molecular (High) | \$118.51 | \$ 117.60 | \$ 115.85 |
| **Cost of Carb NS PSA/ACB (Carba 5)** |  |  |  |
| Suggested Carb NS PSA-ACB (%) | 12% | 20% | 40% |
| ASTar Rapid MALDI + Reflex LFA w/o Syndromic Molecular (Low) | \$4.60 | \$ 7.00 | \$ 13.00 |
| AS*Tar* Rapid MALDI + Reflex LFA w/o Syndromic Molecular (High) | \$5.80 | \$ 9.00 | \$ 17.00 |
| Cost Savings Over Syndromic Molecular (Low) | \$65.40 | \$ 63.00 | \$ 57.00 |
| Cost Savings Over Syndromic Molecular (High) | \$114.20 | \$ 111.00 | \$ 103.00 |

*\* AS*Tar* cost is not included since is constant*

## Discussion

Bloodstream infections and sepsis remain a significant clinical challenge, in part due to their time-sensitive nature, in which diagnostic delays can prolong empiric therapy and delay optimization of antimicrobial treatment. In this study, we have demonstrated a RUO workflow in which remnant bacterial suspension generated during routine AST on the ASTar System, which would otherwise be discarded, can be utilized for downstream pathogen ID by MALDI-TOF MS and for detection of enzymes associated with target resistance markers using LFA.

This remnant material supported robust performance across both applications. All 119 samples tested yielded high-confidence MALDI scores (>2.0), including a small subset of five samples that initially scored below the threshold but achieved high-confidence scoring upon repeat testing. In parallel, LFA results demonstrated high concordance with reference genotype data, including full agreement for CTX-M detection and high agreement for target carbapenemase (KPC, NDM, VIM, IMP, and OXA-48-like) detection using the NG CARBA-5 in Enterobacterales, *P. aeruginosa*, and *A. baumannii*. A single discordant OXA-48 result in Enterobacterales was resolved upon repeat testing. Together, these findings indicate that remnant material from the ASTar system could be suitable for reliable downstream applications despite being an unintended by-product that is typically discarded.

Beyond analytical performance, the value of this approach lies in its impact on laboratory workflows and operational bottlenecks. While rapid AST systems have been shown to shorten the time to actionable AST results, their overall benefit may be constrained by dependence on upstream protocols or instruments, including timely organism ID. By repurposing remnant material, this RUO approach combined early pathogen ID and targeted resistance detection within an existing AST run without introducing additional sample preparation steps. MALDI-TOF MS ID and LFA testing can be initiated well before completion of the full AST report with minimal (<10 min) hands-on time. This approach may accelerate the availability of pathogen information within the laboratory workflow while reducing the need for duplicate sample processing.

This could have important implications for laboratory workflows: The combination of early pathogen ID and targeted resistance marker detection, alongside full rapid phenotypic AST, may support earlier time to results^10,11^. Taken together, this has the potential to enable a simplified workflow when ASTar is involved and is a practical complement to molecular methods rather than a direct replacement, potentially resulting in cost savings compared to syndromic molecular systems.

From an operational perspective, this approach also addresses several key barriers. System cost remains a major barrier to the adoption of diagnostic instruments, and this RUO approach avoids introducing additional high-complexity instrumentation into the workflow^12^. It concurrently reduces labour-intensive protocols by offering a direct-from-blood-culture MALDI-TOF MS without subculturing, while reducing interference from blood-culture matrix components.

Furthermore, automated processing with ASTar ensures reproducible generation of remnant material, minimizes variability associated with manual extraction and is advantageous in laboratories with limited staffing. Flexibility in processing time was also observed, as LFA performance remained unchanged whether pellet preparation and testing were initiated after cartridge availability in ASTar after sample preparation or after refrigeration of the cartridge for six hours, indicating no performance degradation over time. Consistent performance was also observed with reflex LFA results (refrigerated for six hours after retrieval of the cartridge) with the RUO Acineto-5 assay.

Several study limitations should be considered when interpreting these findings. Firstly, this work was conducted under RUO conditions and does not represent a cleared or intended use of the ASTar System. Secondly, the evaluation used contrived PBCs comprising Gram-negative strains with established genotypic reference data, which may not capture the variability encountered in routine clinical samples. Thirdly, organism scope was limited to Gram-negative bacteria, which limited the LFA resistance mechanisms evaluated. Lastly, further studies are needed to evaluate this RUO protocol in real-world clinical settings, including comparison to standard of care methods in routine clinical use, to evaluate its impact on clinical outcomes such as time to effective therapy and de-escalation rates, and their associated outcomes such as I.V.-to-oral switch, and associated patient outcomes, as well as its performance in settings with varying operational schedules.

Notably, MALDI-TOF MS and LFA testing represent two examples of applications of this purified remnant bacterial suspension. While beyond the scope of the present study, this material may support additional downstream uses depending on specific laboratory needs.

In conclusion, this RUO evaluation demonstrates that ASTar remnant bacterial suspension, which is otherwise discarded in the routine rapid AST workflow, has the potential to be utilized for MALDI-TOF MS pathogen identification and LFA-based resistance detection. Consolidating phenotypic AST in approximately six hours with same-run resistance detection or LFA reflex testing reduces reliance on multiple systems and sample preparation steps and may accelerate time to results, offering a cost-effective and operationally efficient alternative to existing approaches.

## Acknowledgements

We gratefully acknowledge the contributions of Dr. Jenny Göransson.

## Abbreviations

BC: blood culture
GN: Gram-negative
BSI: bloodstream infection
AMR: antimicrobial resistance
ID: identification
QC: quality control
SOC: standard of care
AST: antimicrobial susceptibility testing
ICU: intensive care unit
CLSI: Clinical & Laboratory Standards Institute
EUCAST: European Committee on Antimicrobial Susceptibility Testing
BMD: broth microdilution
MIC: minimum inhibitory concentration
PBC: positive blood culture
IVDR: *In Vitro* Diagnostic Regulation
EA: essential agreement
MIN: minor discrepancy
MAJ: major discrepancy
VMJ: very major discrepancy
MALDI-TOF MS: matrix-assisted laser desorption ionization time of flight mass spectrometry

## Author contributions

Conceptualisation and methodology: VG, MM, NB, JG

Validation: IN, SV, ER

Investigation: IN, SV, ER

Analysis: IN, SV, ER

Writing (original draft): OM, VG

Writing, reviewing and editing: All

## Ethics and informed consent

This study does not include factors necessitating patient consent.

## Data availability statement

The datasets used and/or analysed during the current study are available from the corresponding author on reasonable request.

